# *Col4a2-eGFP* mouse model reveals the molecular and functional dynamics of basement membrane remodelling in hair follicle morphogenesis

**DOI:** 10.1101/2023.10.31.564866

**Authors:** Duligengaowa Wuergezhen, Eleonore Gindroz, Ritsuko Morita, Kei Hashimoto, Takaya Abe, Hiroshi Kiyonari, Hironobu Fujiwara

**Author notes:** Ritsuko Morita’s present address: Graduate School of Frontier Biosciences, Osaka University, Suita 565-0871, Japan.

## Abstract

The precisely controlled remodelling of the basement membrane (BM) is considered vital for morphogenesis. However, the molecular and tissue-level dynamics of the BM during morphogenesis and their functional significance remain largely unknown, especially in mammals, due to limited visualization tools. We developed knock-in mouse lines in which the endogenous collagen IV gene (*Col4a2*) was fused with a fluorescent tag. Through live imaging of developing hair follicles, we revealed a spatial gradient in the turnover rate of COL4A2 that is closely coupled with the BM expansion rate. The proliferation of epithelial progenitors coincided with the increased expansion of their underlying BM. Epithelial progenitors displaced with directionally expanding BM, but did not actively migrate on stable BM. The addition of a matrix metalloproteinase inhibitor delayed the turnover of COL4A2, restrained the expansion of the BM, and induced a directional shift in the division angle of epithelial progenitors, altering the hair follicle morphology. Our findings revealed spatially distinct BM dynamics within the continuous epithelial BM and affirmed their significance in orchestrating the proliferation, movement and fate of progenitor cells, as well as the macro-level shape of organs during development.

## Introduction

Multicellular organisms are intricate composites of cells and extracellular matrices (ECMs). ECMs are complex polymer networks of proteins and polysaccharides that provide cells with diverse biochemical and biomechanical cues. One major type of ECM is the basement membrane (BM), which is a thin, dense, sheet-like ECM that surrounds most tissues in metazoans ^1^. The BM contains a core set of proteins, including laminin, collagen IV, nidogen and perlecan, and many other cell- and tissue-specific BM proteins, BM-modifying proteins and morphogens ^1,2^. The BM is an evolutionally ancient ECM that is found across metazoans and is the earliest ECM structure to emerge during development ^3–5^.

The BM plays essential roles in animal development and homeostasis by acting as a versatile solid-phase cell adhesive substrate and signalling platform. By interacting with cells via integrins and other cell surface receptors, this sheet-like ECM regulates various fundamental cell behaviours, including cell adhesion, migration, proliferation, differentiation, apoptosis, polarity and shape. Moreover, compositions of BM proteins undergo spatiotemporal specialization during development ^6–8^. Therefore, the BM dynamically changes its physicochemical properties and organization to orchestrate the fates and behaviours of cells. Deficiencies and mutations in BM genes cause various developmental and homeostatic disorders in multicellular organisms, highlighting the BM’s critical role in the formation and maintenance of multicellular systems ^9–11^.

The BM has long been considered a static support structure, like a floor in a building. However, recent studies suggest that the BM is far more dynamic than previously thought at both the molecular and structural levels ^2,4,12,13^. For instance, the molecular turnover of collagen IV is observed during the development of *Drosophila* and *C. elegans* and appears to contribute to proper organ development ^14–16^. Furthermore, the assembly of laminin and collagen IV may affect the BM’s mechanical properties, influencing the architecture of epithelial tissues during tumour progression ^17^. Therefore, perfectly balanced BM remodelling and turnover, which may control spatial and temporal changes in the BM’s biochemical and biomechanical properties, would be essential for its dynamic functions. However, the molecular and tissue-level dynamics of the BM, along with their underlying regulatory mechanisms and physiological functions, remain largely unexplored.

A significant challenge in BM biology is examining the cross-scale dynamics of varied, dense, complex supramolecular matrices *in vivo*, especially in mammals. The primary approach to visualizing BM dynamics involves genetically tagging fluorescent molecules to BM proteins. However, certain properties associated with the BM can impede the insertion of a large fluorescent protein without affecting normal BM functions. These properties include 1) large modular structures, 2) specialized intracellular transport and secretory mechanisms, 3) post-translational processing, 4) the assembly of supramolecular complexes, 5) complex molecular interactions and 6) unique extracellular physicochemical environments, such as redox status. While a few individuals of *C. elegans* and *Drosophila* have been manipulated to express fluorescently tagged endogenous BM proteins with normal functions ^14,18^, mice expressing fluorescent tag–fused core BM proteins have exhibited functional abnormalities ^19-^ _21_. Therefore, generating mice that express endogenous core BM proteins fused with a fluorescent tag while maintaining normal function remains a major challenge.

The BM comprises two independent, self-assembling networks of laminin and collagen IV that are interconnected by nidogen and perlecan ^22^. Collagen IV networks are crucial for providing BMs’ core structure and tensile strength ^4,23–26^. Collagen IV consists of up to six genetically unique α-chains, which are designated α1(IV) to α6(IV). Each collagen IV polypeptide comprises three distinct domains: a cysteine-rich N-terminal 7S domain of ∼150 amino acids; a central, long, triple-helical collagenous domain of ∼1300 amino acids; and a globular C-terminal NC1 domain of ∼230 amino acids. Heterotrimeric collagen IV molecules can interact through their 7S domains to form a tetramer and vital disulfide bonds between collagen molecules, or through their NC1 domains to form a dimer and create supramolecular network organizations ^16,22,27–29^. Due to their intricate heterotrimeric protein structure and complex intermolecular interactions, various dysfunctional mutations have been reported throughout collagen IV genes, leading to the definition of a broad spectrum of disorders, including embryonic lethality, myopathy, glaucoma, haemorrhagic stroke, nephropathy and cochlear dysfunctions ^23,30,31^. Therefore, alterations to the collagen IV amino acid sequence or the introduction of a large fluorescent protein at any position can potentially impede the synthesis, assembly, deposition and function of collagen IV molecules, posing challenges for fluorescent tagging. Although generating mice that express endogenously tagged collagen IV with normal function would be a powerful approach to visualizing and investigating BM dynamics in mammalian tissues in health and disease, it has not yet been achieved.

In this study, we generated knock-in mouse lines that express fluorescently tagged endogenous collagen IV alpha 2 chains (COL4A2). The mice grew normally and were fertile even when homozygous. Using embryonic skin tissues from these mice, we established a 3D live-imaging method to enable the long-term continuous monitoring of COL4A2 within tissues. Furthermore, our study revealed the remarkably dynamic and spatially distinct turnover of collagen IV proteins, which play a crucial role in the expansion of the BM at the tissue level, the orientation of epithelial cell division and epithelial morphogenesis.

## Results

### Development of a 4D imaging method to visualize BM dynamics in live tissues

To visualize BM dynamics in real time, we first sought to develop knock-in mice that expressed fluorescently tagged endogenous collagen IV. Of the six polypeptide chains of collagen IV, we selected the α2 chain (*Col4a2*) that occurs in the BM of all tissues as the α1α1α2 heterotrimer ^28^. By referring to the insertion site of the GFP protein in the viking^G454^ *Drosophila* collagen IV GFP-trap line ^32^, a cDNA encoding eGFP protein with short linker sequences was inserted into the 7S domain of the collagen IV α2 chain with the CRISPR/Cas9-assisted knock-in method in mouse zygotes ^33^ (Fig. 1A and Fig. S1A). We confirmed successful transgene insertion into the *Col4a2* gene in *Col4a2-eGFP* knock-in pups with PCR screening and genomic sequencing (Fig. S1B and S1C). Importantly, the *Col4a2-eGFP* allele was transmitted at the expected Mendelian ratio of 1:2:1 when knock-in heterozygous mice were bred (Fig. 1B and Table 1). Furthermore, adult homogeneous knock-in mice were visually indistinguishable from their wild-type littermates (Fig. 1C). These results indicated that heterozygous and homozygous *Col4a2-eGFP* knock-in mice developed normally and were fertile.

**Figure 1.**
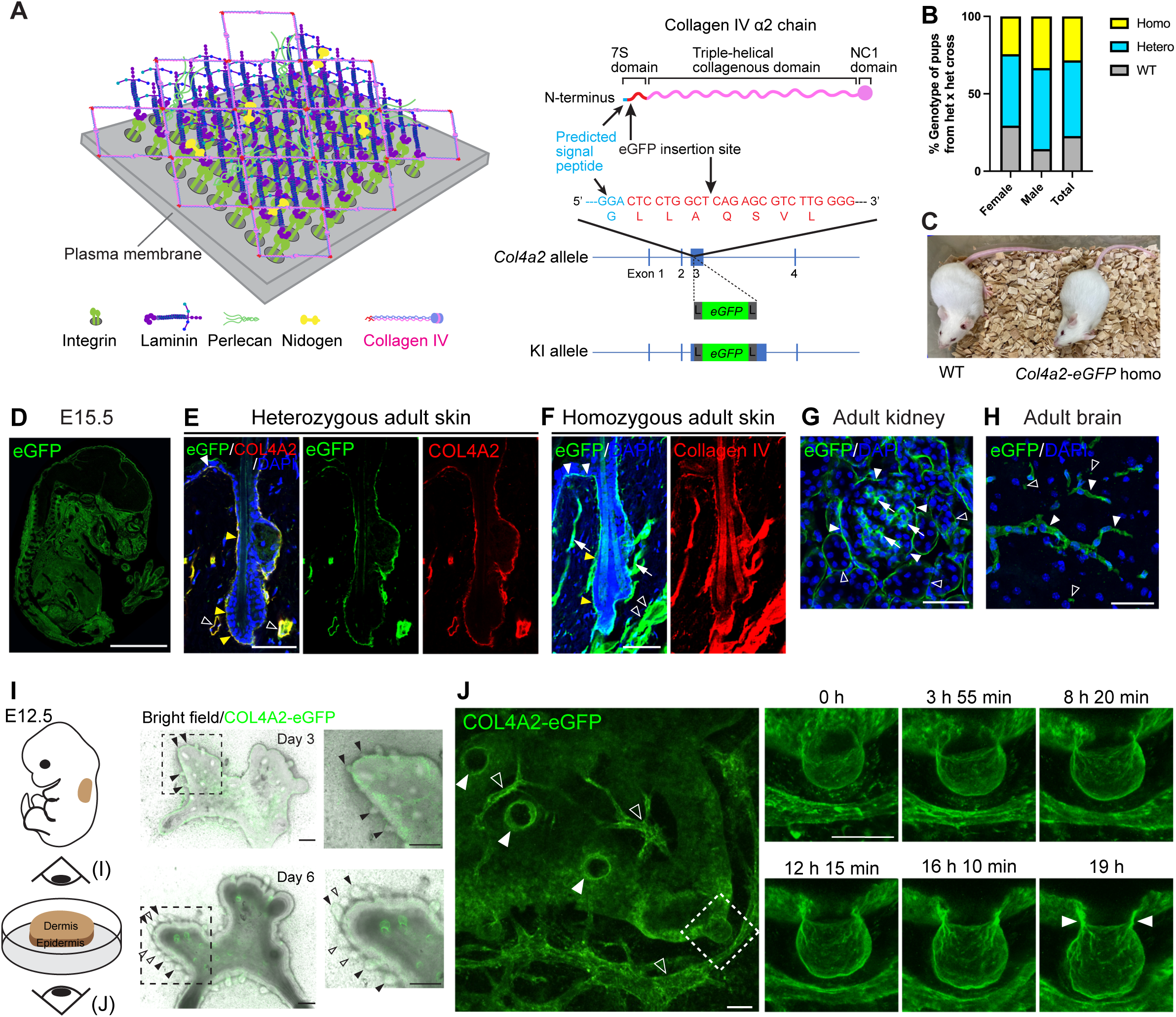
Development of the 4D imaging method for visualizing basement membrane dynamics. A. A schematic model of the BM’s molecular structure and the insertion site for a fluorescent protein in the collagen IV α2 chain. B. Genotypes of pups from *Col4a2-eGFP* het × *Col4a2-eGFP* het crosses. The sample size is reported in Table 1. C. Appearances of 8-week-old wild-type and *Col4a2-eGFP* homo mice. D. Representative immunofluorescence image of E15.5 embryos of *Col4a2-eGFP* mice stained for eGFP. Scale bar: 3 mm. E–H. Representative immunofluorescence images of adult tissues from *Col4a2-eGFP* mice. (E) In P56 adult telogen dorsal skin tissues from het mice, eGFP (green) colocalizes with COL4A2 (red) in the BMs of the epidermis (closed white arrowheads), hair follicle epithelium (closed yellow arrowheads) and vessel-like tissues (open arrowheads). (F) In P56 adult telogen dorsal skin tissues from homo mice, eGFP (green) is distributed as in het mice and colocalizes with collagen IV (red) in the BMs of the epidermis (closed white arrowheads), hair follicle epithelium (closed yellow arrowheads), vessel-like tissues (open arrowheads), and arrector pili muscle (arrows). (G) In adult kidneys, eGFP (green) was detected in Bouman’s capsules (closed arrowheads), the mesangial matrix (arrows) and collecting ducts (open arrowheads). (H) eGFP (green) localizes in capillaries (closed arrowheads) and fractone-like structures (open arrowheads). DAPI (blue) was used for nuclear counterstaining. Scale bars: 50 μm. I. Representative fluorescent stereo-microscope images of cultured dorsal skin explants from E12.5 embryos of *Col4a2-eGFP* mice. Magnified images of the dotted rectangular areas are shown. Closed arrowheads indicate primary hair follicles and open arrowheads indicate secondary hair follicles. Scale bars: 200 μm. J. Representative snapshot images of the 3D maximum projection of cultured skin explants from E12.5 embryos of *Col4a2-eGFP* mice. Closed arrowheads indicate ring-like accumulations of the COL4A2-eGFP signal at the neck regions of hair follicles. Hair follicles, indicated by closed arrowheads in the left panel, grow vertically from the plane of observation. A dashed rectangle indicates a hair follicle growing horizontally from the curled surface of the epithelium. Snapshot images of the 3D time-lapse maximum projection of this hair follicle are shown in the right panels. Open arrowheads indicate vessel-like structures. Scale bars: 50 μm.

**Table 1.** Genotypes of pups from *Col4a2-eGFP* het ξ *Col4a2-eGFP* het crosses.

| Genotype | WT | Heterozygous | Homozygous | Total |
| --- | --- | --- | --- | --- |
| Number of pups born<br>(female:male) | 32 (23:9) | 69 (36:33) | 40 (19:21) | 141 (78:63) |
| Ratio of genotype<br>occurrence | 22.7% | 48.9% | 28.4% |  |

To examine whether the fluorescent fusion protein was appropriately incorporated into the BMs, we immunohistochemically detected eGFP in the embryonic and adult tissues of *Col4a2-eGFP* knock-in mice. eGFP was detected throughout the embryos at E13.5, E15.5 and E17.5, evidenced by a BM-like linear staining pattern at the tissue borders of many organs, including the skin, kidneys and brain (Fig. 1D and Fig. S1D-G). In the adult skin of *Col4a2-eGFP* heterozygous mice, eGFP colocalized with COL4A2 and perlecan in the BMs of epidermal, hair follicle epithelial, arrector pili muscle and vessel-like tissues (Fig. 1E and Fig. S1H), suggesting that the eGFP tags were fused with COL4A2 proteins and that the fusion proteins localized correctly in the BMs. In homozygous mice, antibodies against eGFP, pan-collagen IV and perlecan showed BM-like staining patterns and colocalized with each other (Fig. 1F and Fig. S1I), suggesting that normal BMs were formed even when both endogenous *Col4α2* alleles were replaced by the knock-in alleles. In adult kidneys, eGFP was detected in the glomeruli along the Bowman’s capsule and collecting ducts and in an amorphous pattern in the mesangial matrix (Fig. 1G and Fig. S1J), as stated in a previous report ^34^. In the adult brain, eGFP overlapped with perlecan and localized in the blood vessels, where the blood–brain barrier forms and in fractone-like structures (Fig. 1H and Fig. S1K). We concluded that the COL4A2-eGFP fusion protein was appropriately incorporated into the BMs and functioned normally.

We then examined whether the COL4A2-eGFP fusion protein had a fluorescent intensity that would be sufficient to visualize BM dynamics in long-term live imaging of 3D tissues. We employed an *ex vivo* explant culture system of embryonic dorsal skin that recapitulated the development of hair follicles *in vivo* (Fig. 1I) ^35^. Confocal 3D live imaging captured spatiotemporal changes in COL4A2-eGFP. For example, changes in the shape of BMs were captured in the developing hair follicles and vessels without significant photobleaching (Fig. 1I, J and Supplementary Video 1). Moreover, a ring-like accumulation of COL4A2-eGFP was observed in the neck region of the developing hair follicles (Fig. 1J, closed arrowheads). These results demonstrate that knock-in mice expressing endogenously tagged fluorescent COL4A2 are a viable approach to studying BM dynamics *in situ*.

### Spatially distinct BM expansion rates synchronise with directional cell movement

To investigate how BMs spatiotemporally expand in concert with elongating and shape-changing developing organs, we first bleached eGFP fluorescence in several rectangular regions of the BM to compartmentalize BM zones and measured the changes in the length of these BM compartments (Fig. 2A, B and C). After 7 h of culturing, the BM showed an ∼30% increase in length at the tip of the growing hair follicle, and the rate of elongation gradually decreased towards the upper part of the hair follicle, with no or reduced elongation in the upper stalk region (Fig. 2B and 2C). Therefore, the BM of developing hair follicles does not expand uniformly, like a passive balloon inflating, but rather undergoes spatially distinct expansion, generating a spatial gradient of expansion rates.

**Figure 2.**
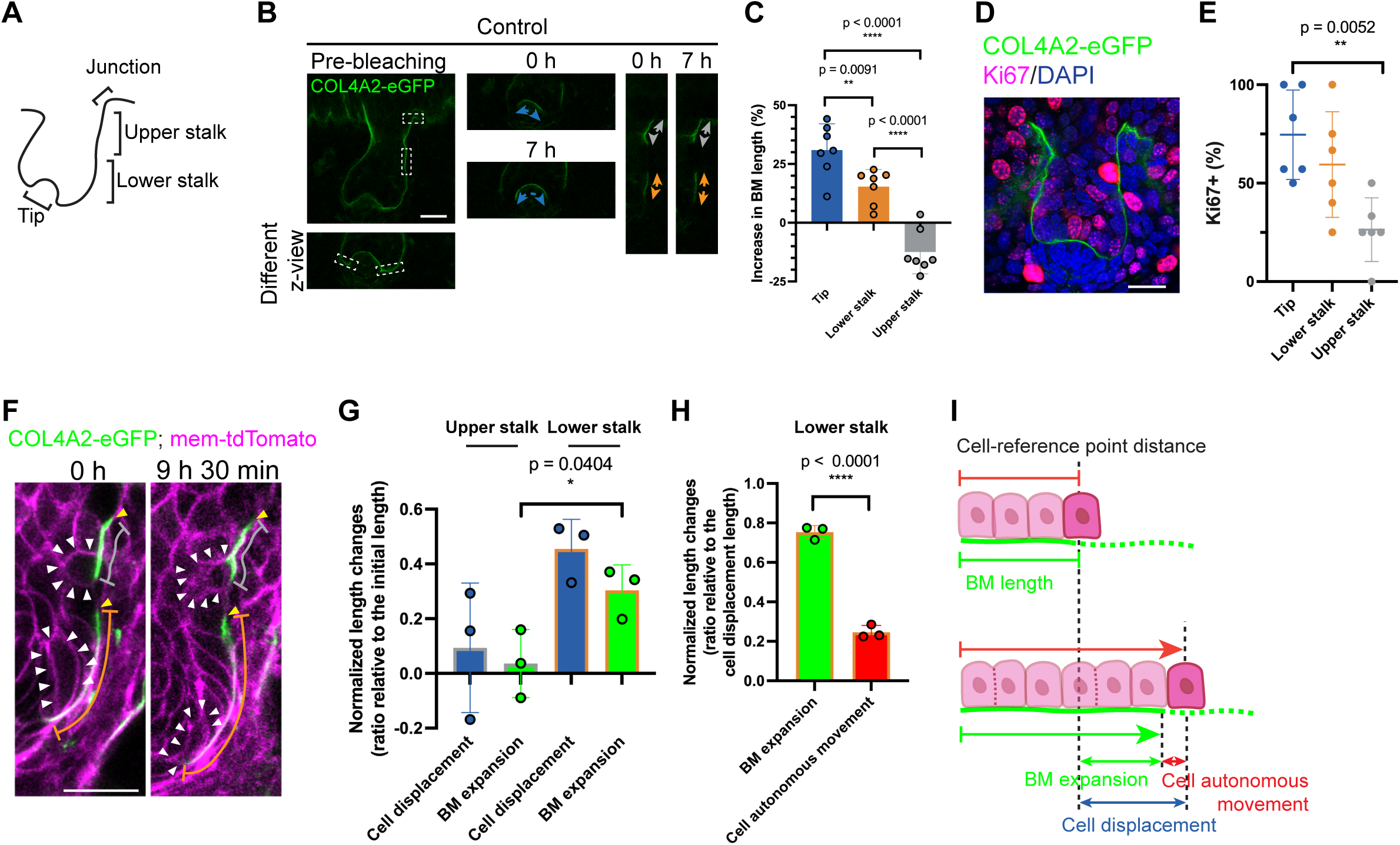
Spatially distinct BM expansion rates synchronise with directional cell movement. A. Schematic diagram of region definitions within the developing mouse hair follicle. Tip: The bending interface region between the pre-matrix and dermal condensate. Lower stalk: The lower half region of the hair follicle extending to the tip region. Upper stalk: The upper half region extending to the junction region. Junction: The bending neck region where the follicular epidermis meets the interfollicular epidermis. Molecular and tissue dynamics of the BM were measured within these distinct BM compartment regions. B. Representative confocal images of hair follicles in dorsal skin explants from *Col4a2-eGFP* mice used to measure changes in BM length. Dotted rectangles indicate the regions selected for photobleaching. BM length changes were measured at the tip (blue lines), lower stalk (orange lines), and upper stalk (grey lines) of the hair follicle during a 7-h culture period. Scale bar: 20 μm. C. Quantification of per cent BM length changes over a 7-h culture period (n = 7 hair follicles, from 4 independent explant/experiment). Values are presented as mean ± SD. Two-tailed unpaired t-tests were used. D. Representative immunofluorescence image of developing hair follicles in embryonic skin explants from *Col4a2-eGFP* mice, stained for eGFP (green) and Ki67 (magenta). DAPI (blue) was used for nuclear counterstaining. Scale bar: 20 μm. E. Quantification of per cent Ki67-positive nuclei in the tip, lower stalk, and upper stalk regions (n = 6 hair follicles from 4 explants/experiments). Values are presented as mean ± SD. A two-tailed unpaired t-test was used. F. Representative snapshot images of 3D time-lapse videos of developing hair follicles from embryonic skin explants of *Col4a2-eGFP; mem-tdTomato* mice. The relative location of the BM (lower bleached edge) and attached epithelial basal progenitors (indicated by white arrowheads) from the reference points on the BM (upper bleached edge, indicated by yellow arrowheads) in both the lower stalk region (orange lines) and upper stalk region (grey lines) were measured at 0 h and 9 h 30 min. Scale bar: 20 μm. G. Bar graph of the normalized length changes (ratio relative to the initial length) in the distance of basal progenitors from reference points on the BM (cell displacement), the distance of bleached edges of the BM from reference points on the BM (BM expansion) (n = 3 cells, each from an independent hair follicle from an independent explant/experiment). Values are presented as mean ± SD. H. Bar graph of the normalized length changes (ratio relative to the cell displacement length) of BM expansion and calculated cell autonomous movement (n = 3 cells, each from an independent hair follicle from an independent explant/experiment). Values are presented as mean ± SD. I. A schematic representation of the composite motion of cells and the BM.

BMs can regulate tissue shape by guiding changes in cellular behaviour or creating patterned physical constrictions ^4^. We next examined the relationships between the BM expansion rate and the proliferation of basal epithelial progenitors. Immunostaining for proliferation marker Ki67 showed that ∼75% of basal epithelial progenitors were proliferative in the tip region, where high BM expansion was observed, and this percentage gradually decreased towards the upper part of the hair follicle. Only ∼25% were Ki67-positive in the upper stalk region (Fig. 2D and 2E), suggesting that the rate of cell proliferation is associated with the degree of BM expansion.

We further examined cellular movements on this expanding BM and considered how much of the observed cell movement could be explained as active-cell autonomous migration and how much was passive displacement due to changes in the BM structure. To track the movement of epithelial basal progenitors and their relative position to the underlying BM, we bleached COL4A2-eGFP fluorescence in several regions of the BM and measured changes in both cell position and length of the BM (Fig. 2F). Over 9.5 hours, cells in the upper stalk region were displaced ∼9% towards the tip of the hair follicle from the reference point on the BM (bleached upper edge), while cells in the lower stalk were displaced ∼46% (Fig. 2G). The underlying BM elongated ∼4% in the upper stalk and ∼30% in the lower stalk towards the tip of the hair follicle. To determine the contribution of cell autonomous movement, including the effects of cell proliferation, to the overall cell displacement, we subtracted the contribution of BM expansion from the cell displacement (Fig. 2H). In the lower stalk region, the contribution of cell autonomous movement was ∼25% of the total cell displacement, while BM expansion contribute to ∼75%. These findings indicate a coordinated directional movement of both cells and the BM towards the tip of the hair follicle. Our measurement results further support this by showing that the bleached edge of the BM moved alongside the cells (Fig. 2F). Altogether, these results suggest that basal epithelial cells move in unison with the directionally expanding underlying BM but do not actively migrate on the stable BM (Fig. 2I).

### Increased BM expansion is coupled with increased turnover of collagen IV protein

The molecular mechanism that facilitates the expansion of preexisting, edgeless BM is largely unknown. It may be driven by the incorporation of new collagen IV protein through homeostatic remodelling of the existing collagen IV network. However, how much incorporation and turnover of the BM molecules occurs spatiotemporally during morphogenesis had not been quantified at the cellular level. Therefore, we measured the dynamics of COL4A2-eGFP fluorescence recovery after photobleaching (FRAP) as a proxy for the incorporation of collagen IV protein into the BM (Fig. 3A and Supplementary Video 2). Surprisingly, unlike the slow turnover rates of collagens reported in adult animals (from days to months) ^36–38^, the fluorescence at the tip region of the hair follicle recovered by 50% just 2 h 40 min after bleaching, indicating a remarkably rapid COL4A2 incorporation rate (Fig. 3A and 3B). Fluorescence recovery was evenly observed across the large bleached area but did not occur from its edges, suggesting that the recovery of fluorescence occurred through the incorporation of COL4A2-eGFP from outside the BM rather than via diffusion from adjacent unbleached BM areas. In contrast, the COL4A2-eGFP of the lower stalk and upper stalk regions recovered by about ∼15% and ∼8%, respectively, indicating a relatively slow COL4A2 incorporation rate (Fig. 3B). Furthermore, to monitor the dynamics of the COL4A2 protein present at specific locations and timepoints, we created a knock-in mouse line named *Col4a2-mKikGR*, which expresses endogenous COL4A2 fused with the photoconvertible fluorescent protein mKikGR. Photoconverted red COL4A2-mKikGR in the BM of the lower stalk region gradually decreased and almost completely disappeared during 8–10 hours of culturing, while green fluorescence gradually increased and replaced the red signal (Fig. 3C and Supplementary Video 3). This red-to-green colour change indicated the turnover of the COL4A2 protein, where pre-existing COL4A2-mKikGR proteins were replaced by newly synthesized or recruited proteins. In contrast, in the upper stalk and the follicular-interfollicular junction regions, photoconverted red COL4A2-mKikGR remained. These results suggest a rapid COL4A2 turnover rate at the tips of developing hair follicles. We also observed that the red fluorescence did not diffuse from the photoconverted region. Therefore, the regional BM expansion rate is tightly linked to the regional turnover rate of COL4A2.

**Figure 3.**
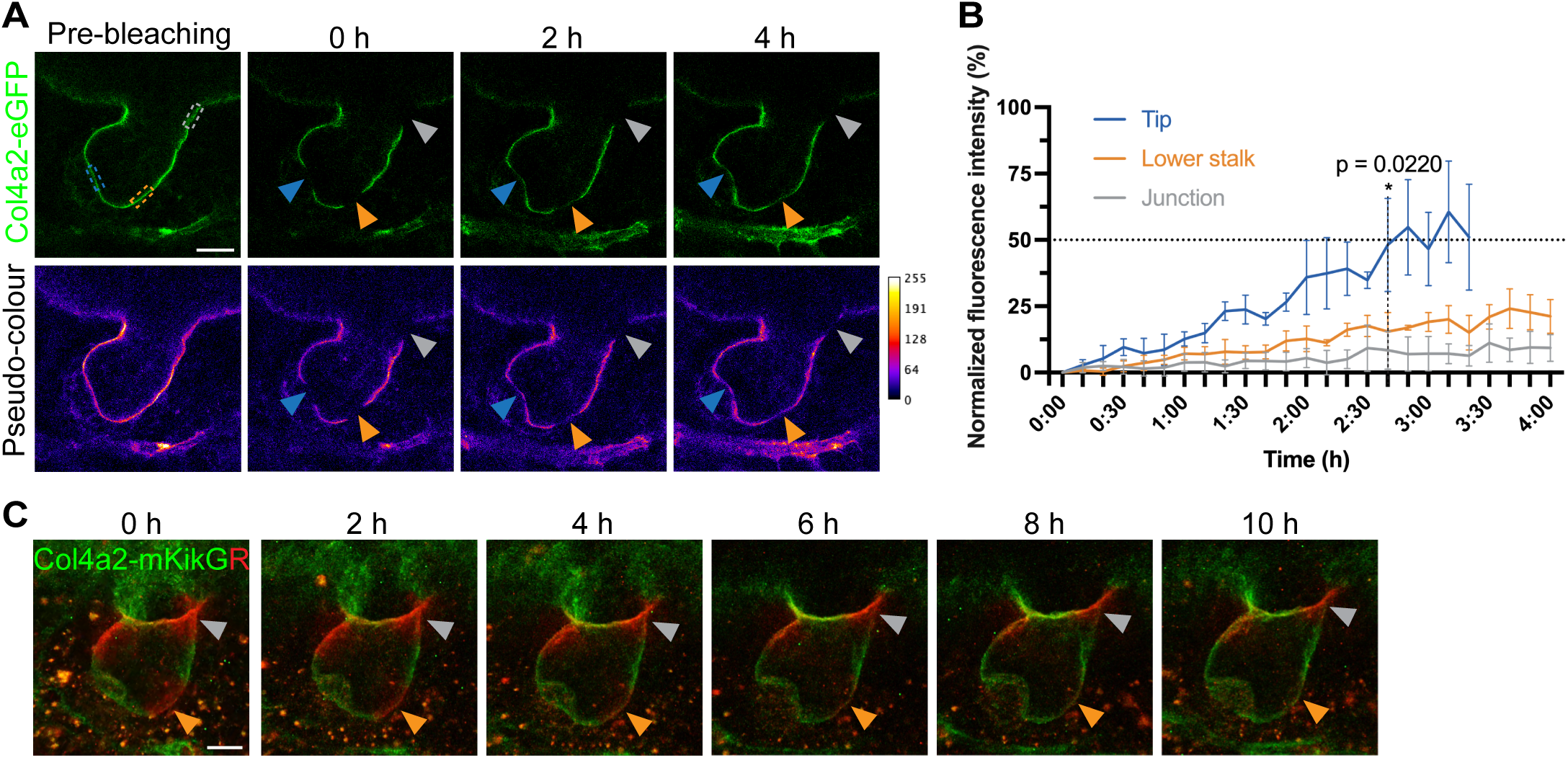
Spatially distinct collagen IV turnover. A. Representative time-lapse images of COL4A2-eGFP FRAP experiments. Confocal single-colour images (upper) and pseudo-colour images (lower) show the BM of a developing hair follicle in explants that were photobleached in the tip (blue arrowheads), lower stalk (orange arrowheads) and follicular–interfollicular junction (grey arrowheads) regions before and after photobleaching at selected recovery times. Scale bar: 20 μm. B. Line graph of the normalized fluorescence recovery of COL4A2-eGFP over 4 h (n = 3 hair follicles, each from an independent explant/experiment). Values are presented as mean ± SD. A two-tailed unpaired t-test was used. C. Representative maximum projection images of COL4A2-mKikGR in developing hair follicles in embryonic skin explants from *Col4a2-mKikGR* mice at the indicated times after photoconversion. Scale bar: 20 μm.

### Matrix metalloproteinases are required for COL4A2 incorporation, BM expansion and hair follicle morphogenesis

The proteolytic cleavage of ECM components plays essential roles in ECM remodelling ^9^. At the molecular level, proteolysis of the BM components may nick the assembled BM proteins, which may stimulate their degradation and metabolism, rendering the BM more pliable and creating new insertion sites for free BM proteins. This process can facilitate the metabolism or turnover and remodelling of the BM. The central enzymes for ECM remodelling are matrix metalloproteinases (MMPs) ^9^. Recent advancements in ECM imaging have begun to reveal that protease activity in MMPs is required for BM molecular and tissue-level dynamics, as well as effective branching morphogenesis and early embryonic development _14,15,39_. Therefore, we hypothesized that MMP activity is required to generate the observed BM dynamics in hair follicle morphogenesis at the molecular and tissue levels. We tested this by treating skin explants with a broad-spectrum peptidomimetic MMP inhibitor, batimastat (BB-94), which binds to the active sites of MMPs and inhibits the enzymatic activities of several MMPs, including MMP-2 and MMP-9, which cleave collagen IV ^40^. With this inhibitor, the fluorescence recovery of COL4A2-eGFP in the FRAP assays was greatly delayed in all regions of hair follicle BMs (Fig. 4A, 4B and Supplementary Video 4). The fluorescence in the tip region of the control hair follicles recovered by 50% just 2 h 40 min after bleaching, while the inhibited samples recovered by only ∼5%. Similarly, the lower stalk and junction regions showed drastic reductions in recovery rate. These quantitative data indicate that the enzymatic activities of MMPs are crucial for COL4A2 incorporation into BMs.

**Figure 4.**
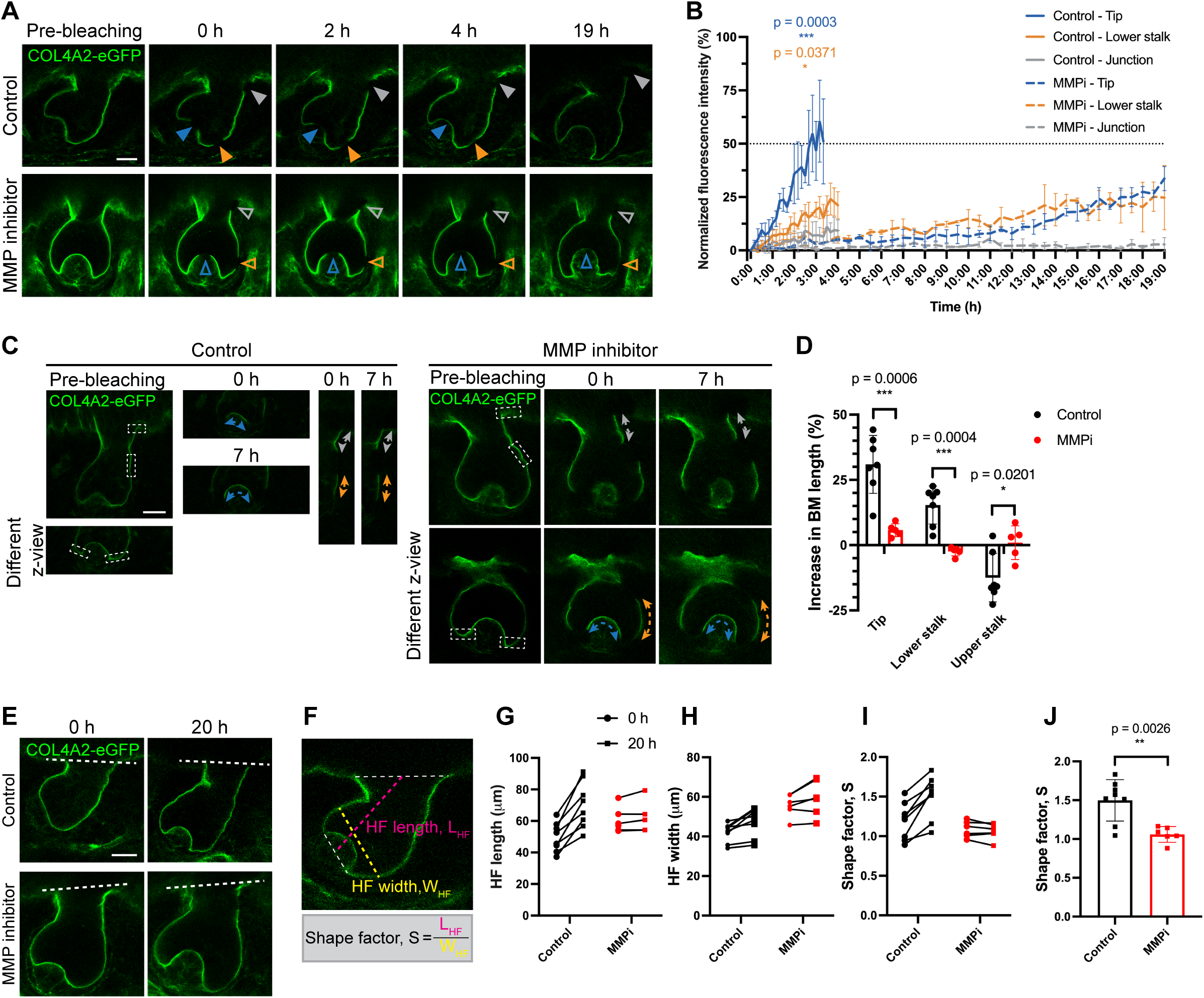
Matrix metalloproteinases are required for COL4A2 incorporation, BM expansion and hair follicle morphogenesis. A. Representative snapshot images of COL4A2-eGFP FRAP experiments with developing hair follicles in skin explants from *Col4a2-eGFP* mouse embryos with or without an MMP inhibitor. Images show the different recovery rates of COL4A2-eGFP (green) after photobleaching the tip (blue arrowheads), apex (orange arrowheads) and junction (grey arrowheads) regions of control and 3 μM batimastat-treated (MMP-inhibited) follicles (open arrowheads). Scale bar: 20 μm. B. Quantification of the normalized mean intensity values of COL4A2-eGFP fluorescence in the bleached regions of FRAP experiments in (A) (n = 3 hair follicles, each from an independent explant/experiment). MMPi indicates MMP inhibitor treatment. Values are presented as mean ± SD. Two-tailed unpaired t-tests were used. Control data of this experiment are shown in Figure 3B. C. Representative snapshot images to measure BM expansion in developing hair follicles in skin explants from *Col4a2-eGFP* mouse embryos. COL4A2-eGFP fluorescence in rectangular regions that were photobleached. The BM length change at the tip (blue lines), lower stalk (orange lines), and upper stalk (grey lines) in control or MMP inhibitor-treated samples was measured. Control data from this experiment are shown in Figure 2B. Scale bar: 20 μm. D. Quantification of the changes in BM length in the experiments depicted in (C) (n = 7 hair follicles from 4 independent explants/experiments for the control, and n = 5 hair follicles from 3 independent explants/experiments for the inhibitor-treated condition). Values are presented as mean ± SD. Two-tailed unpaired t-tests were used. Control data of this experiment are shown in Figure 2C. E. Representative confocal images of control and MMP inhibitor-treated developing hair follicles at 0 h and 20 h. Some data were obtained from the hair follicles used in the FRAP analysis. Scale bar: 20 μm. F–J. Quantitative analysis of the shape of the developing hair follicles cultured under control or MMP-inhibited conditions. (F) Representative image of the measurement of the shape factor S. L_HF_ indicates the length of the hair follicles and W_HF_ indicates the width of the hair follicles. The shape factor, S, was determined by dividing the L_HF_ by the W_HF_. Graphs show the changes in the length of hair follicles (G), width of hair follicles (H) and shape factor (I) during a 20-h culture period. (J) Graph of the rate of changes in S during the 20-h culture period (n = 8 hair follicles from 7 independent explants/experiments for the control, and n = 6 hair follicles from 3 independent explants/experiments for the inhibitor-treated condition). Values are presented as mean ± SD. A two-tailed unpaired t-test was used.

We then investigated changes in the BM expansion rate in MMP inhibitor–treated hair follicles. Quantitative analysis demonstrated that, with the MMP inhibitor, the tip region only showed a ∼6% increase over a 7-h culture period, not the ∼30% increase observed in the control (Fig. 4C and 4D). There was almost no elongation in other regions. These results indicate that enzymatic activities of MMPs are required for COL4A2 incorporation and BM expansion. Moreover, the differences in collagen IV protein turnover dynamics are closely linked to the tissue-level expansion of BMs, generating the spatial gradient in BM expansion rate by regulating the local COL4A2 turnover rate.

Alterations in BM dynamics and structure could significantly affect organ morphogenesis and shape ^17,41^. Therefore, we examined organ shape changes in developing hair follicles from the hair germ to hair peg stages with and without the MMP inhibitor. Eighteen hours after the addition of the MMP inhibitor, we started live imaging observation of the shape changes of hair follicle epithelia, using the COL4A2-eGFP signal to define the basal outline of the hair follicle epithelia. Hair follicles exhibited an abnormal, disproportionate shape when treated with an MMP inhibitor (Fig. 4E). Inhibitor-treated hair follicles showed halted hair follicle elongation (denoted as L_HF_; Fig. 4F and 4G) and increased hair follicle width (denoted as W_HF_; Fig. 4F and 4H). We describe these distinct tissue architectures as a shape factor, S, defined as the ratio of L_HF_ to W_HF_. We found high and increasing S values in control hair follicles, indicating the formation of an elongating cylindrical structure under normal development (Fig. 4I). In contrast, inhibitor-treated hair follicles showed non-increasing, low S values, indicating the formation of widening hair follicle structure compared to the control (Fig. 4I and 4J). We conclude that MMPs’ enzymatic activities are required to incorporate and turnover COL4A2 proteins, as well as to expand the BM. The inhibition of these activities disrupts hair follicle morphogenesis.

### Inhibiting matrix metalloproteinases alters the orientation of the daughter cell allocation of epithelial progenitors

The spatiotemporal coordination of cell proliferation and spatial arrangement of cells is one of the major determinants of organ shape ^42^. To interrogate the potential causes of the wider shape of developing hair follicles after the administration of the MMP inhibitor, we examined the cell division pattern of epithelial basal progenitors in live images (Fig. 5A). During a 9.5-h imaging period, ∼25% of basal cells divided in control follicles, while in inhibitor-treated hair follicles, this rate decreased to ∼15% (Fig. 5B). We also performed EdU incorporation assays; at the tips of control follicles, ∼60% of basal cells were positive for EdU, a percentage that gradually decreased towards the upper part of the hair follicle (Fig. 5C and 5D). In inhibitor-treated hair follicles, EdU-positive basal cells decreased in the tip and lower stalk regions, while no notable decrease was observed in the upper stalk region, where the effects of the MMP inhibitor on BM dynamics were small (Fig. 5C and 5D). These data indicate that MMP inhibition reduced the proliferation of basal cells, through which the BM is dynamically remodelled and expanded.

**Figure 5.**
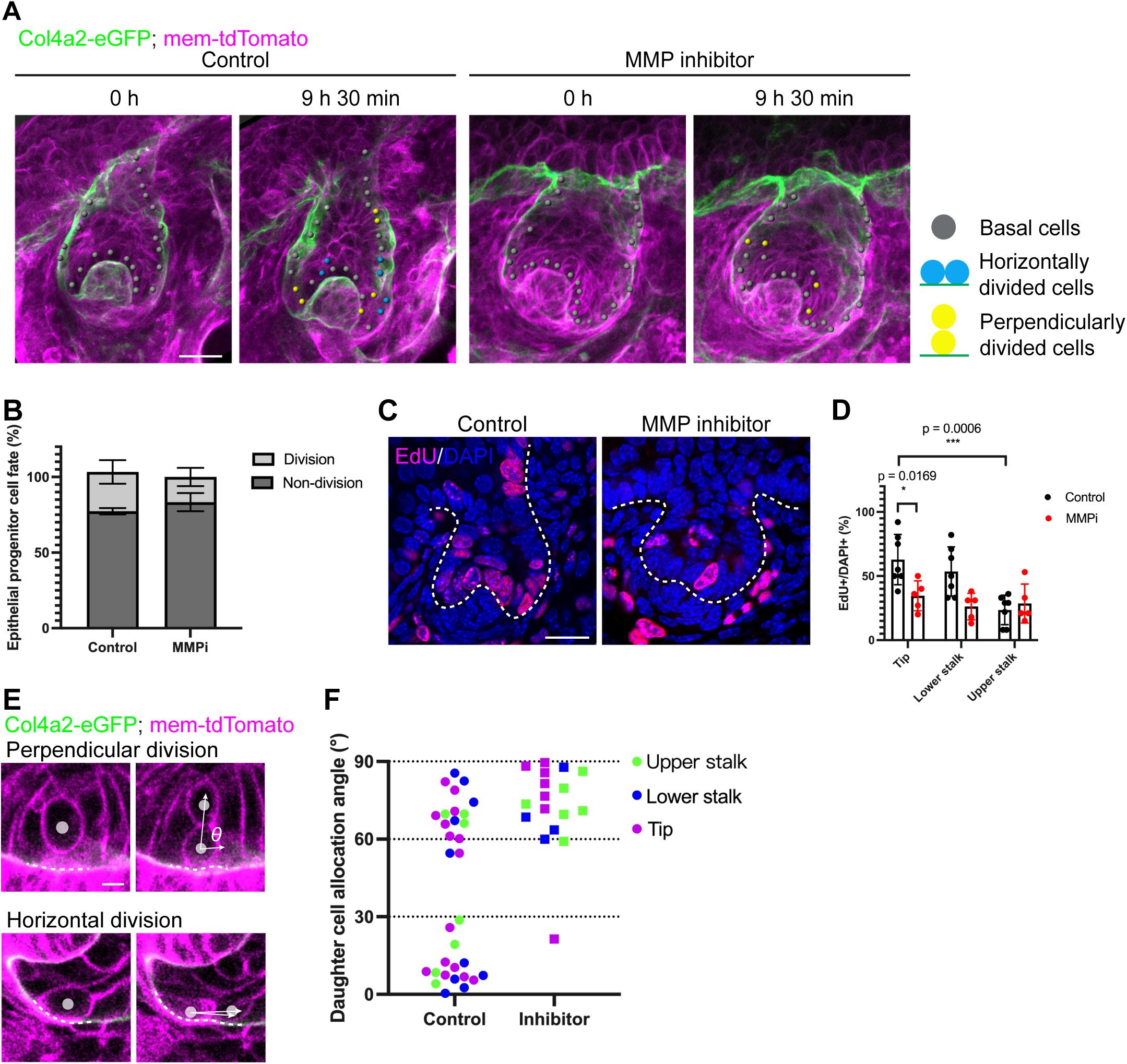
Matrix metalloproteinase inhibition alters the angle of daughter cell allocation. A. Representative snapshot images and cell division tracking data from developing hair follicles in skin explants from *Col4a2-eGFP; mem-tdTomato* mouse embryos. The BM and cell membrane were visualized with COL4A2-eGFP (green) and mem-tdTomato (magenta), respectively. Scale bars: 20 μm. B. Quantification of the cell division of epithelial basal progenitors in the experiments in (A) (n = 3 hair follicles, each from an independent explant/experiment). C. Representative results of the EdU incorporation assays. Dividing cells were labelled with EdU (magenta). All cells were counterstained with DAPI (blue). Scale bar: 20 μm. D. Quantification of the per cent of EdU-positive basal epithelial cells in the tip, lower stalk and upper stalk regions (n = 7 hair follicles from 3 independent explants/experiments for the control, and n = 5 hair follicles from 3 independent explants/experiments for the inhibitor-treated condition). Values are presented as mean ± SD. Two-tailed unpaired t-tests were used. E. Daughter cell allocation angle of proliferating basal epithelial progenitors relative to the BM zone in the experiments in (A). Representative images of perpendicular and horizontal divisions are shown. Scale bar: 5 μm. F. Plots illustrating the angle of daughter cell allocation relative to the BM under control and MMP inhibitor treatment conditions (n = 32 cells from 7 hair follicles from 5 independent explants/experiments for the control, and n = 17 cells from 5 hair follicles from 5 independent explants/experiments for the inhibitor-treated condition.

We used our live imaging data to further examine the angle of postmitotic daughter cell allocation compared to the BM. In the control hair follicles, 50% of divided cells formed cell allocation angles lower than 30% (8 < 30°; horizontal) and the remaining 50% formed cell allocation angles of more than 60% (8 > 60°; perpendicular) (Fig. 5A, 5E and 5F and Supplementary Video 5). Strikingly, with the MMP inhibitor, ∼94% of proliferated daughter cells were allocated to perpendicular angles (suprabasal direction). Therefore, the orientation of the daughter cell allocation of basal epithelial progenitors is tightly linked to MMP-dependent COL4A2 protein dynamics and BM expansion. These results suggest that the increased cell supply towards the centre of epithelial tissue in the tip and lower stalk regions contributed to generating the wider hair follicle tissue shape observed under the inhibition of MMP activity.

## Discussion

In this study, we successfully generated knock-in mice that express fluorescence-tagged endogenous COL4A2. Through live imaging and the quantitative analysis of COL4A2 dynamics in hair follicle development, we revealed a notable spatial gradient in the COL4A2 turnover rate and the BM expansion rate (Fig. 6). Importantly, these two processes exhibit a tightly coupled relationship. Furthermore, epithelial progenitors move together with the expanding BM, and the inhibition of MMPs substantially suppresses both COL4A2 turnover and BM expansion. This inhibition completely shifts the division angle of epithelial progenitors towards the perpendicular. Meanwhile, epithelial tube elongation halts, and a wider tube shape emerges. Our findings elucidate the interplay between COL4A2 turnover, BM expansion and epithelial progenitor behaviour, illuminating the complex orchestration of tissue morphogenesis via the molecular and functional dynamics of the BM.

**Figure 6.**
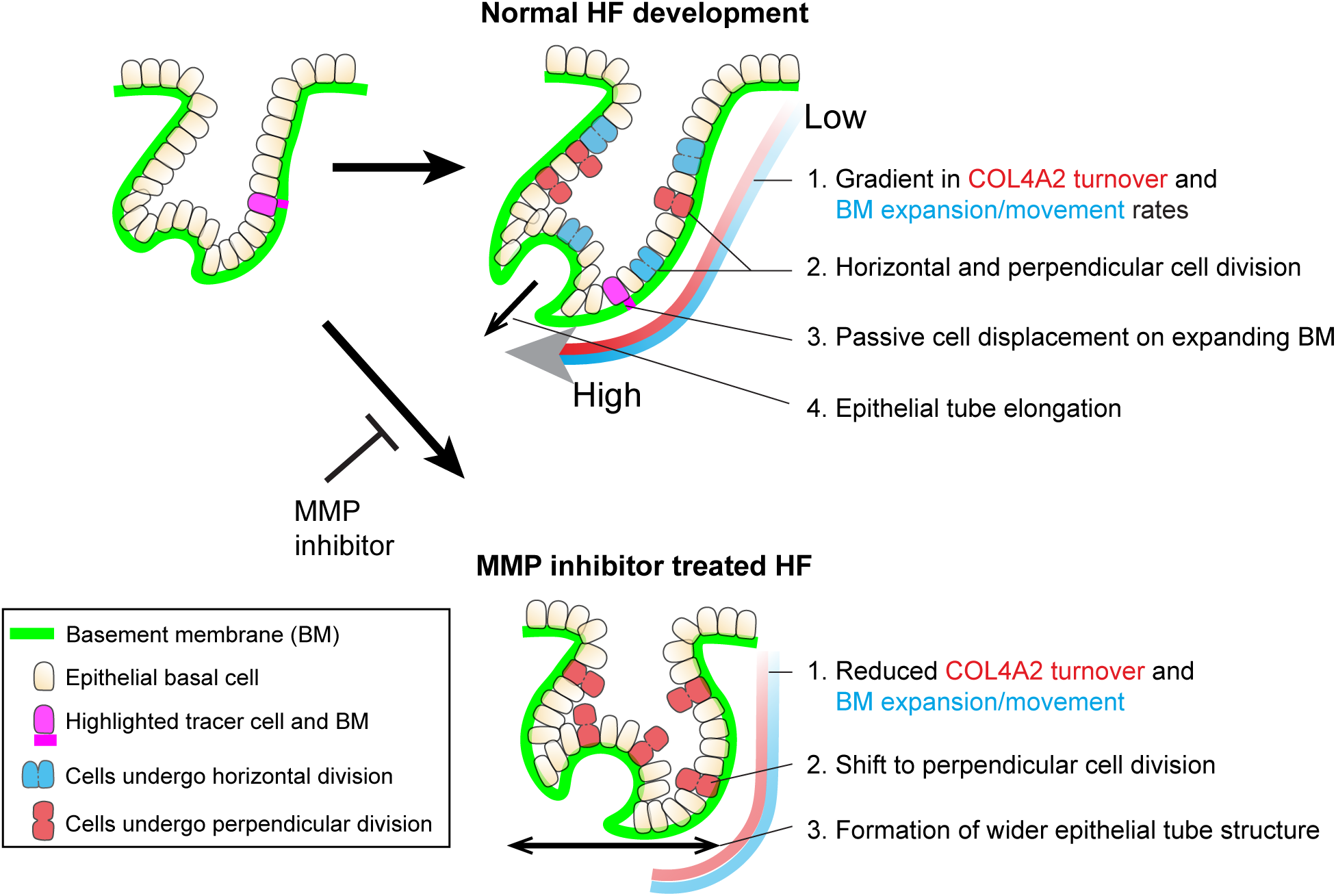
Model depicting the role of BM dynamics in hair follicle morphogenesis. This model illustration summarizes the spatially distinct molecular and tissue-level dynamics of the BM, highlighting its pivotal role in regulating cell and tissue dynamics during hair follicle morphogenesis. Green lines indicate BMs. Magenta cells and BM zones are highlighted tracer epithelial basal cells and BM zones, respectively. Blue cells are epithelial progenitors dividing horizontally relative to the BM, and red cells are epithelial progenitors dividing perpendicularly to the BM. In normal hair follicle development, the following key features are observed: (1) the rates of COL4A2 turnover and BM expansion exhibit spatial gradients; (2) epithelial progenitors undergo both horizontal and perpendicular divisions; (3) cells undergo passive displacement on the directionally expanding BM rather than actively migrating on a stable and immobile BM; and (4) epithelial tube results in directional elongation. In MMP inhibitor treated hair follicles: (1) COL4A2 turnover and BM expansion are restrained; (2) progenitors alter the angle of daughter cell allocation from horizontal to perpendicular; (3) directional epithelial elongation is halted, resulting in a wider organ shape.

### The challenges and importance of fluorescence tagging of endogenous BM proteins

A major challenge in ECM biology is quantitatively examining the dynamics of supramolecular assembled complex matrices *in situ* at both the molecular and tissue levels. Traditionally, fluorescence-labelled antibodies to ECM proteins have been used to visualize ECMs in tissues, yielding valuable insights into ECM dynamics during morphogenesis ^39,43,44^. However, antibody labelling methods have certain limitations, including the possible masking of active epitopes, steric hindrance, the variable efficiency of ECM protein labelling across time and space and the selective binding of antibodies to specific forms of the target ECM proteins ^45,46^. Another frequently used method is the injection of fluorescently labelled ECM proteins ^17,47^. However, this approach may also have some limitations, as the observed dynamics, locations and functions could vary with the proximity, availability and biological activity of the added protein. Thus, to more accurately quantify ECM dynamics, the expression of endogenously tagged and functionally normal fluorescent fusion ECM proteins is required.

Previously, several successful cases of altered core BM proteins had been reported in *C. elegans* ^14,48,49^ and *Drosophila* ^18^, but very few in mammals. This difficulty may stem from the intricate supramolecular assembly process that BM molecules undergo and their extensive interactions with cell surface receptors, other ECM molecules and soluble factors ^50,51^. Therefore, the introduction of a large fluorescent protein into BM proteins may cause undesirable consequences, such as the misfolding or misassembling of the fusion proteins ^52^ or inhibition of their functions and interactions with other proteins ^43^. In *C. elegans*, mNeonGreen was inserted at the N-terminus of the collagen IV α2/*let-2* gene, resulting in viable heterozygotes but nonviable homozygotes ^14^. There was an attempt to adapt the Dendra2 tagging of laminin β1 used in *C. elegans* ^53^ to mammalian cells, but secretion failed _19_. More recently, a knock-in mouse model that expresses endogenously tagged laminin β1-Dendra2 fusion protein has been developed and used to identify BM-producing origins and dynamics in breast tumours ^21^. In this mouse model, crossing heterozygous mice produces homozygous mice only 1% of the time, and they exhibit syndactyly and testis dysplasia. Another mouse line, expressing mCherry-tagged human nidogen-1 under the Rosa26-CAG promoter, has been generated and used to label BMs, but unclear BM signals and non-BM-like signals were also observed in several tissues ^20^. These studies underscore the difficulties in generating mice that express fluorescently tagged, functionally normal BM proteins.

In this study, we inserted short linker-tagged eGFP or mKikGR near the N-terminus of the 7S domain of the endogenous *Col4a2* gene, avoiding the putative N-terminus intermolecular covalent cross-linking sites and the collagenous triple-helical domain of the 7S domain ^32,54,55^. These results suggest that although 7S domains are intermolecularly cross-linked and play essential roles in collagen IV network formation and stability, their N-terminus ends might have structural tolerance for the insertion of exogenous molecules. This site can be used to tag them with other fluorescent proteins, bioactive molecules and biosensors, such as morphogens, pH sensors and calcium sensors, in BMs *in vivo*.

### Cells and BMs move together

Embryonic development involves large-scale cell movement and tissue deformation. The dynamics of cell behaviour during this process have been primarily attributed to the coordinated interplay of active cell shape changes, cell migration and cell proliferation, while the ECM has been considered a passive, stable substrate for these cell-autonomous activities _13,42,56,57_. In mouse hair follicles, important morphogenetic cell behaviours, such as increased cell proliferation and directional epithelial cell movement towards the leading edge, occur during development and regeneration ^35,58^. In addition to these cellular dynamics, our BM live imaging and quantitative analysis revealed an intriguing phenomenon: the BM exhibits spatial gradients of expansion rates, resulting in directional expansion along the axis of organ elongation. Moreover, epithelial progenitors proliferate and move with the directionally expanding BM rather than actively migrating on a stable and immobile BM. When BM expansion is limited by MMP inhibitors, the progenitors alter the angle of daughter cell allocation from horizontal to perpendicular, halting the directional organ elongation and resulting in a wider organ. These findings suggest that directional BM expansion contributes to the coordinated and directed expansion of tubular epithelia during morphogenesis by regulating the angle of daughter cell allocation and cell displacement. Tissue stretching can induce planar cell division in epithelial cells ^59–61^. Furthermore, intrinsic tissue-scale BM motion can displace adhering cells ^62^. Therefore, BM expansion and movement may also actively influence cell division and the migration of epithelial tissue in other organs.

BM movement has been observed in other systems via different methodologies. For instance, in avian embryos, vortex-like movement of fibronectin-containing sub-epiblastic ECM, which appears to contain both the BM and the underlying interstitial matrix, has been reported ^47^. Another study of *Hydra* demonstrated that both the epithelia and the mesoglea, which comprise the BM and fibrillar matrix, moved together towards the tips of feet and tentacles ^63^. Furthermore, during salivary gland branching morphogenesis, directional movement of the collagen IV network has been observed ^39^. Interestingly, in this context, the collagen IV network moves from the tips to the stems of the branches, the opposite of the direction of BM movement in developing hair follicles, while cell motion during the development of salivary glands remains undefined. Therefore, the quantitative measurement of composite tissue motion, encompassing both cellular and matrix components, will greatly enhance our understanding of the active roles of BM dynamics in various biological phenomena, including morphogenesis, tissue regeneration and disease progression.

### Mechanisms and roles of the spatial gradient in the turnover ratio of COL4A2

What causes the spatial gradients in BM expansion rates? We show that the spatial gradient in the turnover rate of COL4A2 corresponds well with the rate of BM expansion. Furthermore, the inhibition of MMP activity strongly hampers both COL4A2 turnover and BM expansion, suggesting that the extent of MMP-mediated BM protein cleavage and the assimilation of new BM proteins into the matrix determine the degree of BM expansion. In this scenario, the spatial organization of MMP activity would be crucial for regulating BM dynamics and various epithelial morphogenic events, such as invagination, tube extension and branching ^64^. In hair follicles, the expression patterns and roles of MMPs are largely unknown. However, MMPs have been relatively well-studied in the mammary gland: MMP2-null mammary glands are deficient in terminal end bud invagination but have excessive secondary branching, whereas MMP3-null mammary glands exhibit normal invagination but have deficient secondary branching ^65^. Furthermore, the broad-spectrum MMP inhibitor BB-94 halts BM movement and perforation and the epithelial branching of the salivary gland ^39^. MMP inhibition in early post-implantation mouse embryos also prevents BM perforation and inhibits primitive-streak extension ^66^. Therefore, BM dynamics should be investigated across different scales, from the molecular to the tissue level, to better understand the functional importance of BM dynamics for various biological events.

How does MMP cleavage of BM proteins promote BM turnover and expansion? The proteolysis of collagen IV can create new cut surfaces within the collagen IV network, allowing the incorporation of free collagen IV molecules into the BM. We show that treatment with MMP inhibitors slows the incorporation of COL4A2 into the BM and halts BM expansion, supporting the notion that the proteolysis of the BM components makes the BM pliable and creates new sites for the insertion of BM molecules, enhancing the molecular turnover, remodelling and expansion of the BM ^13^. Notably, the different properties of the BM are not independent; rather, they are intertwined such that changes in one BM feature often coincide with changes in other aspects ^9^. For example, in addition to affecting BM turnover and expansion, MMP-mediated collagen IV cleavage may render the collagen IV network more flexible and softer, making it more susceptible to shape changes. Indeed, the gradient of BM stiffness contributes to directed tissue elongation in *Drosophila* egg chamber development ^26^. Therefore, the MMP-mediated molecular-level dynamics of the BM could influence organ shape by regulating the various physical, biochemical and biomechanical properties of BMs.

In summary, the BM live-imaging approach established in this study offers potential for studying the role of BM dynamics in a wide range of biological phenomena in mammals, including early development, organogenesis, homeostasis, regeneration and disease progression for conditions such as cancer. Therefore, this imaging technique makes a significant contribution to our quest for a deeper understanding of the functional importance of BM dynamics in multicellular systems as composites of cells and ECMs.

## Supplementary materials

**Figure S1.**
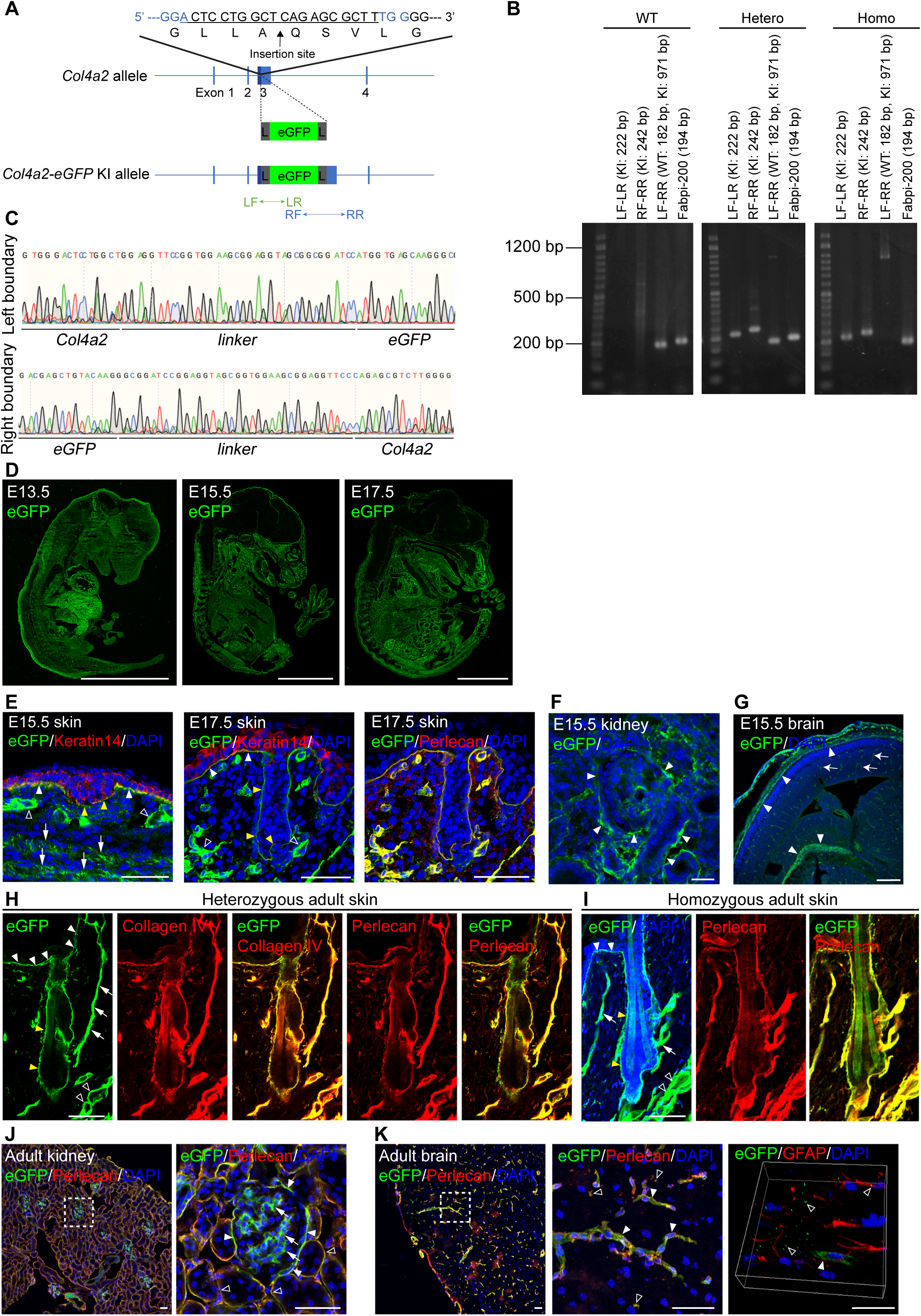
Generation of *Col4a2-eGFP* knock-in mice. A. Targeting strategy for generating *Col4a2-eGFP* knock-in mice. An eGFP protein was inserted into the N-terminus region of the 7S domain of the COL4A2 protein (between A^28^ and Q^29^). To minimize the structural interference of the inserted fluorescent protein with COL4A2 protein structure and functions, a glycine- and serine-rich flexible linker, GGSGGSGGSGGS, was added to both ends of the EGFP protein. L, linker; LF, left forward primer; LR, left reverse primer; RF, right forward primer; RR, right reverse primer. B. Representative image of PCR screening to distinguish wild-type, heterozygous and homozygous *Col4a2-eGFP* mice. Fabpi-200: An internal control primer pair for detecting *Fabpi* gene. 50 bp DNA ladder RTU (GeneDirex) is used. C. Genome sequences of boundaries between the *Col4a2* gene and linkers. D. Representative immunofluorescence images of E13.5, E15.5 and E17.5 embryonic tissues of *Col4a2-eGFP* mice stained for eGFP. Scale bars: 3 mm. E–G. Representative immunofluorescence images of tissues from *Col4a2-eGFP* mouse embryos. (E) At E15.5 (left panel), the eGFP (green) signal is detected at the epidermal BM (white closed arrowheads) below the keratin 14-positive (red) basal epidermis, the BMs of the hair placode (yellow closed arrowheads), the dermal vessel-like structures (open arrowheads) and the panniculus carnosus muscle-like layers (arrows). At E17.5 (middle and right panels), eGFP (green) is detected in the perlecan-positive (red) epidermal and vessel-like BMs. (F) In the E15.5 kidney, eGFP (green) appears around the tubular epithelial structures (closed arrowheads). (G) In the E15.5 brain, eGFP (green) appears in pial BM zones (closed arrowheads) and vessel capillaries (arrows). DAPI (blue) was used for nuclear counterstaining. Scale bars: (E, F) 20 μm, (G) 200 μm. H–K. Representative immunofluorescence images of adult tissues from *Col4a2-eGFP* mice. (H) In P56 adult telogen dorsal skin tissues from het mice, eGFP (green) colocalizes with collagen IV (red) and perlecan (red) in the BMs of the epidermis (closed white arrowheads), hair follicle epithelium (closed yellow arrowheads), arrector pili muscle (arrows) and vessel-like tissues (open arrowheads). (I) In P56 adult telogen dorsal skin tissues from homo mice, eGFP (green) is distributed as in het mice and colocalizes with perlecan (red). (J) In adult kidneys, eGFP (green) is detected in Bouman’s capsules (closed arrowheads), the mesangial matrix (arrows) and collecting ducts (open arrowheads). (K) In the subventricular zone, eGFP (green) colocalizes with perlecan (red) in capillaries (closed arrowheads) and fractone-like structures (open arrowheads) around the GFAP+ (red) astrocytes. DAPI (blue) was used for nuclear counterstaining. Scale bars: 50 μm.

**Video 1. Representative time-lapse video capturing COL4A2-eGFP signals in mouse embryonic dorsal skin explant cultures**

Representative time-lapse video of the 3D maximum projection of cultured skin explants from E12.5 embryos of *Col4a2-eGFP* mice, corresponding to Fig. 1J. The BM was visualized with COL4A2-eGFP (green). The BMs of hair follicles, interfollicular epidermis and vessels are visualized.

**Video 2. Representative time-lapse video of FRAP experiments with COL4A2-eGFP in developing hair follicles**

COL4A2-eGFP signals (green) were photobleached in designated regions and signal recovery was monitored. Cultured skin explants from E12.5 embryos of *Col4a2-eGFP* mice were used, corresponding to Fig. 3A.

**Video 3. Representative time-lapse video of pulse-chase experiments in the developing hair follicles of skin explants from *Col4a2-mKikGR* mouse embryos**

Green COL4A2-mKikGR signals were photoconverted to red and the changes in these COL4A2-mKikGR signals were chased over about 19 h at 30 min intervals, corresponding to Fig. 3C.

**Video 4. Representative time-lapse video of FRAP experiments with COL4A2-eGFP in developing hair follicles under MMP inhibition**

COL4A2-eGFP signals (green) were photobleached in designated regions and signal recovery was monitored under MMP inhibition with batimastat. Cultured skin explants from E12.5 embryos of *Col4a2-eGFP* mice were used, corresponding to Fig. 4A.

**Video 5. Representative time-lapse video of daughter cells undergoing horizontal or perpendicular division during hair follicle development**

This time-lapse video shows tracked epithelial basal progenitors that undergo horizontal or perpendicular division in cultured skin explants from *Col4α2-eGFP; mem-tdTomato* mouse embryos. The BM and cell membrane were visualized with COL4A2-eGFP (green) and mem-tdTomato (magenta), corresponding to Fig. 5E.

## Materials and Methods

### Mice

All experimental procedures were approved by the Institutional Animal Care and Use Committee of the RIKEN Kobe Branch. The care and handling of the mice adhered to the ethical guidelines of the RIKEN Kobe Branch as well. To visualize BM dynamics, we generated *Col4a2-eGFP* knock-in mice (Accession No. CDB0100E: https://large.riken.jp/distribution/mutant-list.html). A 717-bp eGFP sequence (without a stop codon) was inserted into the three amino acids after the end of the predicted signal peptide (M^1^–G^25^) on exon 3 of *Col4a2*. To minimize the structural interference of the inserted fluorescent proteins with COL4A2 functions, a *Gly*- and *Ser*-rich flexible linker, (GGS)_4_, was added to both ends of the eGFP gene. To make a donor vector that contained homologous arms (319 bp and 286 bp), the *Col4a2* genomic sequence around the eGFP insertion site was cloned into the pBluescript II. The cDNA sequences of eGFP and the linkers were integrated into the insertion site within the donor vector. C57BL/6 zygotes were subjected to microinjection with the mixture of Cas9 protein (100 ng/μl), crRNA (50 ng/μl)/tracrRNA (100 ng/μl) and the donor vector (10 ng/μl), and then the zygotes were transferred to pseudo-pregnant females to obtain founder knock-in mice ^33^. The crRNA (5’-ACU CCU GGC UCA GAG CGU CUG UUU UAG AGC UAU GCU GUU UUG-3’) and the tracrRNA (5’-AAA CAG CAU AGC AAG UUA AAA UAA GGC UAG UCC GUU AUC AAC UUG AAA AAG UGG CAC CGA GUC GGU GCU-3’) were purchased from FASMAC. The knock-in mice were screened via PCR-based genotyping using the three primer pairs (LF+LR, RF+RR, LF+RR) shown in Supplemental Figure 1. The primers were LF (5’-GCTGCTGCTAGCAACTGTGACAG-3’), LR (5’-GTGCAGATGAACTTCAGGGTCAGC- 3’), RF (5’-GGTCCTGCTGGAGTTCGTGACC-3’) and RR (5’-AGCACACACCTACAATGCACACG-3’). The LF and RR primer pair was designed on the *Col4a2* allele and the RF and LR pair was designed on the *eGFP* sequence. Three primer pairs were used for PCR genotyping to distinguish the WT allele (LF-RR; 182 bp); and the *Col4a2-eGFP* knock-in allele (LF-LR; 222 bp, RF-RR; 242 bp; LF-RR; 971 bp) (Supplemental Fig. 1B). We used genome sequencing to confirm that the linker and *eGFP* construct was correctly inserted into the *Col4a2* gene (Supplemental Fig. 1C). To investigate BM turnover, *Col4a2-mKikGR* mice (Accession No. CDB0126E: https://large.riken.jp/distribution/mutant-list.html) were generated using the same strategy. The plasmid for mKikGR was purchased from addgene (# 54656) ^67^. The knock-in mice were screened via PCR-based genotyping using the three primer pairs (LF1+LR1, RF1+RR, LF1+RR). The primers were LF1 (5’- AAGACTGGGATCATGGACCG-3’), LR1 (5’- TGGCTCCCATTTGACGGTCTTCC-3’), RF1 (5’- ACAGTCATAGAGGGCGGACCTCT - 3’) and RR (5’-AGCACACACCTACAATGCACACG-3’). The LF1 and RR primer pair was designed on the *Col4a2* allele and the RF1 and LR1 pair was designed on the *mKikGR* sequence. Three primer pairs were used for PCR genotyping to distinguish the WT allele (LF1-RR; 363 bp); and the *Col4a2-mKikGR* knock-in allele (LF1-LR1; 689 bp, RF1-RR; 751 bp; LF1-RR; 1134 bp). To visualize cells and the BM simultaneously, *Col4a2-eGFP* mice were bred with *mTmG* mice (The Jackson Laboratories, JAX Strain no. 007576). All mice except for *Col4a2-mKikGR* were bred with FVB/NJcl mice (CLEA Japan) to avoid imaging interference from melanin deposition.

### Immunohistochemistry

For fluorescence immunohistochemistry, tissues were embedded in OCT, frozen and cryosectioned (10–16 μm). The sections were fixed with 4% paraformaldehyde (PFA) in phosphate-buffered saline (PBS) for 5 min at 4 °C, blocked and permeabilized with blocking buffer (0.5% skim milk/0.25% fish skin gelatine/0.5% Triton X-100/PBS) for 1 h at room temperature (RT), and then incubated with the primary antibodies overnight at 4 °C. Then, the sections were washed with PBS and incubated with fluorescence-labelled secondary antibodies for 2 h at RT. The sections were then washed with PBS and mounted with DAKO Fluorescent Mounting Medium. Fluorescent signals were captured under a TCS SP8X (Leica) confocal microscope with LAS X software (version [BETA] 3.5.7.23723).

For whole-mount immunostaining, tissues were fixed with 4% PFA in PBS for 5–60 min at 4 °C. The tissues were blocked and permeabilized with blocking buffer (0.5% skim milk/0.25% fish skin gelatine/0.5% Triton X-100/PBS) for 1 h at 4 °C, and then incubated with the primary antibodies overnight at 4 °C. Then, the sections were washed with PBT (0.2% Tween-20 in PBS, pH 7.4) and incubated with fluorescence-labelled secondary antibodies for 2 h at RT. The sections were then washed three times with PBT (0.2% Tween-20 in PBS, pH 7.4) for 30 min, dehydrated in 50% methanol/PBS for 10 min, 100% methanol for 5 min, 100% methanol for 10 min, 50% benzyl alcohol with benzyl benzoate (1:2) (BABB)/methanol for 5 min, 100% BABB for 5 min and 100% BABB for 10 min. Fluorescent signals were captured under a TCS SP8X (Leica) confocal microscope with LAS X software (version [BETA] 3.5.7.23723). The antibodies used in this study are shown in Supplementary Table 1.

### *Ex vivo* culture of embryonic dorsal skin

The *ex vivo* culture of embryonic dorsal skin was conducted as described previously ^35^. The pregnant mice were euthanized by cervical dislocation and embryos at embryonic days (E) 12.5–13.5 were collected. The entire dorsal skins of the embryos were dissected with 25G needles under a stereo microscope, and a piece approximately 1/4 the size of the entire skin was excised for culturing. Approximately 10 µl of collagen type I-A (Nitta Gelatin) gel solution was added to an empty 35-mm Lumox dish (Sarstedt) on ice, and then the skin piece was embedded into the collagen gel with the dermis facing upwards. The skin/collagen gel drop was incubated in a humidified CO_2_ incubator with 5% CO_2_ at 37 °C for 30 min to set the collagen gel, and it was subsequently submerged in 1 ml of DMEM/Ham’s F12 with L-Glutamine and Sodium Pyruvate (Wako) supplemented with 20% foetal bovine serum (GIBCO), 100 units/mL of penicillin and 100 µg/mL of streptomycin (GIBCO, Grand Island, NY, USA), 1X GlutaMAX (GIBCO) and 10mM HEPES (GIBCO) and 100 μg/ml ascorbic acid (Sigma). The explants were cultured at 37 °C in a humidified atmosphere with 5% CO_2_.

To inhibit MMPs’ enzymatic activity, on day 3 of the culture, the culture medium was replaced with the same medium containing 3 μM of batimastat (Abcam; ab142087). All of the subsequent experiments were performed with a minimum incubation period of 16 h after the addition of the MMP inhibitor.

### EdU incorporation assays

On day 4 of the culture, 10 μM of EdU was introduced into the culture medium of the skin explants using a 10 mM stock solution from the Click-iT™ Plus EdU Cell Proliferation Kit (Thermo Fisher Scientific; C10640). For the MMP inhibitor-treated group, EdU was added 16 h after the addition of the batimastat. After 3 h of incubation with the EdU solution, the skin explants were fixed and permeabilized according to the manufacturer’s protocol. EdU was detected via Click-iT Plus reaction, followed by Hoechst33342 nuclear counterstaining.

### Live imaging of embryonic dorsal hair follicles

After skin tissue dissection, explants were cultured in a CO_2_ incubator for 3 days to induce hair follicle formation. Live imaging of dorsal hair follicles was performed under a TCS SP8X (Leica) confocal microscope with LAS X software (version [BETA] 3.5.7.23723), a stage top incubator (Tokai Hit) and a 25× water-immersion objective lens (Leica, HC FLUOTAR 25×/0.95 W VISIR) or an LSM980 (Carl Zeiss) with a two-photon laser unit Chameleon Discovery NX (Coherent), Zen software (version 2.3), a stage top incubator and a 32× multi-immersion objective lens (Zeiss, C-Achroplan 32×/0.85 W Corr M27). For two-photon laser imaging, a 920 nm laser and a 1040 nm laser were used to excite eGFP and tdTomato, respectively. For long-term live imaging, immersion water was dispensed onto the objective lens using a Water Immersion Micro Dispenser (Leica) or Liquid Dispenser (Merzhauser Wetzlar). Successive optical section images were obtained as stacks at 512 × 512 pixels for the x–y plane, and a z-stack step size of 1–1.5 μm, covering a total tissue depth of ∼30 μm. These image stacks were acquired at intervals of 10–30 min.

### Fluorescence recovery after photobleaching (FRAP)

Photobleaching was performed under a TCS SP8X (Leica) confocal microscope or an LSM980 (Carl Zeiss) with the same accessory settings described above. A 488 nm laser (100% power) was used to induce photobleaching of COL4A2-eGFP in a ROI with a length of 15±2 μm and a width of 2–3 μm. The fluorescence intensity values of the BM region were acquired using ImageJ (version 1.53). First, a line was drawn on the BM of the pre-bleached and bleached regions on three single z-plane images, which included the target z-stack plane and its upper and lower stacks, and fluorescence intensity values were obtained for each pixel. To create line graphs, the fluorescence intensity values of each line from the three z-planes were averaged, and then they were divided by the average fluorescence intensity values of the pre-bleached times.

### Photoconversion

Tissues expressing COL4A2-mKikGR were imaged with a 488 nm laser under a TCS SP8X (Leica) confocal microscope with the same accessory settings described above. A 405 nm diode laser (20% power with five iterations) was used for photoconversion on rectangle ROIs (15 × 15 μm). The photo-converted mKikGR was imaged with a 552 nm laser.

### Measurement of changes in BM length and cell position

To simultaneously analyse changes in BM length and cell position, *Col4a2-eGFP* mice were crossed with *mTmG* mice (JAX stock# 007576). To make reference points for measurement, photobleaching was performed on the BM. To maintain a distinct bleached edge, the initial bleached areas underwent multiple rebleaching sessions during time-lapse imaging. These sessions utilized a 488 nm laser set at 100% power for three sets of 20 iterations each and were conducted every 2–3 h. An imaging interval of 20–30 min for 3D live imaging enabled the consistent identification of the same cells. The length of the BM was measured between the bleached edges of COL4A2-eGFP in the upper stalk and lower stalk regions. Cell positions were determined by measuring from the upper bleached edge of the reference point on the BM to the midpoint of the cell membrane of the target cell in the upper stalk and lower stalk regions.

### Measurement of the angle of epithelial basal cell division

The orientation of epithelial basal cell division was determined by measuring the angle between a line connecting the centres of two daughter cells and the reference line drawn parallel to the underlying BM ^68^. The plasma membrane and the BM were simultaneously visualized with mem-tdTomato and COL4A2-eGFP, respectively. The cells were tracked manually through identification of the cell shape and position, referring to mem-tdTomato at each time point using Imaris (version 8.4.2, Bitplane). Cells undergoing division were identified by observing the following sequence of events: an initial increase in cell volume, mitotic cell rounding, cleavage furrow formation and, finally, the separation of the daughter cells.

### Statistical analysis and reproducibility

No statistical method was used to predetermine the sample size. Statistical parameters, including the numbers of samples and replicates, types of statistical analysis and statistical significance are indicated in the results, figures and figure legends. For the quantitative analysis, we ensured three or more biological replicates for each experiment.

## Data availability

Live imaging data in this study have been stored in the SSBD repository at https://doi.org/10.24631/ssbd.repos.2023.04.298. All data that support the findings of this study are available within the paper and its supplementary information files or from the corresponding author upon reasonable request.

## Acknowledgements

We thank the Laboratory for Animal Resources and Genetic Engineering for their technical assistance; the members of the Fujiwara laboratory for valuable reagents and discussions; Shigeru Kuratani (RIKEN) for sharing animal resources; Yasuko Tomono (Shigei Medical Research Institute) for antibodies against the Collagen IV α2 chain; and Yasunori Miyamoto (Ochanomizu University) for his generous support on K.H.’s visit to the Fujiwara laboratory. This work was funded by RIKEN intramural funding, the RIKEN BDR Stage Transition project, the JSPS Kakenhi (19K22631, 20H03706, 22K19453), the MEXT Kakenhi (23H04928) and the JST CREST (JPMJCR1926) (all to H.F.). D.W. was supported by the RIKEN International Program Associate (IPA) and the Interdisciplinary Program for Biomedical Sciences, Osaka University. K.H. was supported by the Program for Leading Graduate School of Ochanomizu University.

## Author contributions

A. W. performed the experiments and formal analysis and wrote the initial draft. D.W., E.G., K.H., T.A., H.K. and H.F. developed the knock-in mice. R.M. supported the tissue culture experiments. H.F. conceptualized the study, reviewed and edited the initial draft and supervised the project.

